# On statistical tests of functional connectome fingerprinting

**DOI:** 10.1101/443556

**Authors:** Zeyi Wang, Haris Sair, Ciprian Crainiceanu, Martin Lindquist, Bennett A. Landman, Susan Resnick, Joshua T. Vogelstein, Brian Caffo

## Abstract

Fingerprinting of functional connectomes is an increasingly standard measure of reproducibility in functional magnetic resonance imaging connectomics. In such studies, one attempts to match a subject’s first session image with their second, in a blinded fashion, in a group of subjects measured twice. The number or percentage of correct matches is usually reported as a statistic. In this manuscript, we investigate the statistical tests of matching based on exchangeability assumption in the fingerprinting analysis. We show that a nearly universal Poisson(1) approximation applies for different matching schemes. We theoretically investigate the permutation tests and explore the issue that the test is overly sensitive to uninteresting directions in the alternative hypothesis, such as clustering due to familial status or demographics. We perform a numerical study on two functional magnetic resonance imaging (fMRI) resting state datasets, the Human Connectome Project (HCP) and the Baltimore Longitudinal Study of Aging (BLSA). These datasets are instructive, as the HCP includes techinical replications of long scans and includes monozygotic and dyzogotic twins as well as non-twin siblings. In contrast, the BLSA study incorporates more typical length resting state scans in a longitudinal study. Finally, a study of single regional connections is performed on the HCP data.

## 1 Introduction

Fingerprinting of functional connectomes is an increasingly standard measure of reproducibility in functional magnetic resonance imaging connectomics. In such studies, one attempts to match a subject’s first session image with their second, in a blinded fashion, in a group of twice measured subjects. The number or percentage of correct matches is reported as the statistic. In practice, often functional connectivity profiles, correlation matrices from resting state functional magnetic resonance imaging (rs-fMRI) data, are matched. Under such settings, identification accuracies as high as 94% for the Human Connectome Project (HCP) data or as high as 55% for data with more standard quality have been reported (Waller et al., 2017; Finn et al., 2015; Van Essen et al., 2013). The moniker fingerprinting comes from the idea of the fingerprint as a unique person-specific identifier.

Under the hypothesis of exchangeability of the labels, the number or percent of matches is then analyzed relative to a reference permutation distribution to establish evidence of reproducibility, or lack thereof. This area is bolstered by novel applications (Finn et al., 2015; Airan et al., 2016) and large scale replication studies (Zuo et al., 2014; Van Essen et al., 2013) as well as general interest in fMRI reproducibility (Choe et al., 2017; Poldrack and Poline, 2015; Choe et al., 2015; Landman et al., 2011; Griffanti et al., 2016; Shou et al., 2013; Aron et al., 2006). This manuscript details a series of thoughts on the statistical tests associated with matching experiments for the purposes of establishing evidence of reproducibility and includes an analysis using the technique on benchmark datasets.

We summarize our main points as follows: a) the implied, but rarely stated, definition of the null hypothesis for the permutation test is exchangeability of labels; b) this is a weak null which may result in unintended high power for typical alternative hypotheses; c) the null distribution of the permutation test, almost regardless of the permutation strategy, is well approximated by a Poisson(1); d) evidence beyond the test result is desirable for assessing reproducibility; e) covariates can be associated with the matching performance and require further investigation.

It is interesting to note the historic connections of fingerprinting within the field of statistics. None other than statistical luminary Francis Galton was a seminal figure in rigorously establishing fingerprints for identity verification (Stigler, 1995; Caplan, 1990). We do not further discuss connections with Galton’s work, or the century of work on forensic identity verification following, since our fMRI applications only loosely correspond to identity verification as a goal. Instead, the primary concern is the use of matching for establishing the strength of a metric. Nonetheless we continue to use the term “fingerprinting” throughout, as it has been commonly used in the context of functional MRI to denote the ability of imaging to identify a subject, which then implies inherent metric of reproducibility and uniqueness.

## 2 Connectome fingerprinting mechanics

The most common form of matching tries to match one measurement, say the second, to the first. We write our notation out generally, as our thoughts apply broadly beyond that of functional neuroimaging. Let *W_ij_* be the data vector (image measurement) on session *j* = 1, 2 for subject *i* = 1*, …, n*. To perform matching, one requires a distance or similarity metric, *d*(*·, ·*), such as a correlation or inverse correlation over the elements of *W_ij_*. Most applications in fMRI do not demand that *d* formally satisfies the mathematical requirements to be a distance metric, though it is usually symmetric in its arguments. We assume smaller values imply greater similarity.

Let *d_ij_* = *d*(*w_i_*_1_*, w_j_*_2_) be the distance between subject *i* on occasion 1 and subject *j* on occasion 2 given observation *W_ik_* = *w_ik_*, *i* = 1*, · · ·, n*, *k* = 1, 2. Let *m_i_* be the subject label of the best match for subject *i*. Of course, the term “best” is in reference to a matching strategy and we will use *m_i_* generically regardless of which strategy was used. As an example stategy, consider, *m_i_* = argmin*_r_ d_ir_*. Under this scheme, subjects on occasion 2 can be matched multiple times if they are the best match for more than one subject. Because of this, we call this strategy **matching with replacement** (or MWR).

A matrix form is an often preferable method to represent the data. Let *B* be a matrix with a 1 in position *i, j* if subject *i* on sampling occasion 1 is best matched with subject *j* on occasion 2. That is, *B* = [*b_ij_*]_*i,j*_where *b_ij_* = *I*{*m_i_* = *j*} where *I*{*a* = *j*} is an indicator that returns 1 if *a* = *j* and 0 otherwise, *m_i_* is the observed value of *m_i_*. It is interesting to note that matrices of these forms are exactly bootstrap resampling matrices. Table 1 gives an example for *n* = 4. Recall that the first row, (0, 1, 0, 0), implies that among the occasion 2 measurements, subject 2’s is the best match for the occasion 1 measurement of subject 1. The second row, (0, 1, 0, 0), implies subject 2’s occasion 2 measurement is correctly matched to the subject’s occasion 1 measurement. Thus, in this case, subject 2’s occasion 2 measurements are matched twice, for both subject 1 and subject 2 on occasion 1. The standard statistic measurement agreement is the number of correct matches (the trace of *B*, *tr*(*B*)). In our example, the statistic value would be 3.

**Table 1:** Example resampling matrix from matching with replacement. Here the statistic value is 3.

| Time 1 | Time 2 |  |  |  | Total |
| --- | --- | --- | --- | --- | --- |
|  | 1 | 2 | 3 | 4 |  |
| 1 | 0 | 1 | 0 | 0 | 1 |
| 2 | 0 | 1 | 0 | 0 | 1 |
| 3 | 0 | 0 | 1 | 0 | 1 |
| 4 | 0 | 0 | 0 | 1 | 1 |
| Total | 0 | 2 | 1 | 1 | 4 |

Alternatively, one could match **without replacement** (or MWOR). That is, find the best permutation of subjects on the second occasion to match up with the first. As an example, let Γ be the collection of *n ×* 1 vectors of permutations of the integers 1*, …, n*. Then consider

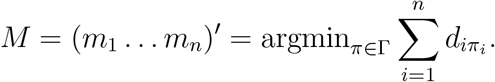

The Hungarian algorithm allows that this optimization can be performed in polynomial time (Pentico, 2007). This is a harder optimization problem, because the optimization is conducted simultaneously and not sequentially, as in the matching with replacement. It is possible to have a non-unique best match. However, given the size and noise of neuroimaging, data the best match is usually unique for the best permutation. If this result is put into a matrix with *b_ij_* = *I*{*m_i_* = *j*}, then *B* is a permutation matrix (a 0,1 matrix with row and column totals all equal to one). Again, the relevant statistic is the trace. Table 2 shows an example with *n* = 4 that has statistic value equal to 2.

**Table 2:** Example resampling matrix from matching without replacement. Here the statistic value is 2.

| Time 1 | Time 2 |  |  |  | Total |
| --- | --- | --- | --- | --- | --- |
|  | 1 | 2 | 3 | 4 |  |
| 1 | 0 | 1 | 0 | 0 | 1 |
| 2 | 1 | 0 | 0 | 0 | 1 |
| 3 | 0 | 0 | 1 | 0 | 1 |
| 4 | 0 | 0 | 0 | 1 | 1 |
| Total | 1 | 1 | 1 | 1 | 4 |

## 3 Inference

Permutation-based inference is the norm in this area. One typically repeatedly permutes the subject labels at occasion 2 and re-performs the matching at each iteration to obtain a null distribution. Given the dimension of the characteristics being matched on, it is typical for no ties to exist in the *d_ij_*, so that the best matches are all unique at each iteration.

This permutation test is motivated by an implicit exchangeability assumption. That is, the underlying null distribution of the statistic is the same for any permutation. Alternatively, the null hypotheses can be developed under stronger iid sampling assumptions.

Despite the apparent simplicity of the permutation procedure, the implementation, hypothesis specification, and inferential interpretation is far from being straightfoward. One of our main results is to show that under nearly all sampling strategies the null distribution of the test statistic is well approximated by a Poisson with a mean of one. The implication of this result is both simple and widespread: the use of the permutation test is unnecessary, as the null hypothesis will be rejected under the same conditions, when *tr*(*B*) is larger than 3 or 4, say, depending on the desired Type I error rate. Thus, computation time and costs can be systematically reduced using this simple, slightly unexpected, but powerful statistical result. Below we provide details on the implicit assumptions associated with the permutation test and the interpretation given these results.

### 3.1 Exchangeability and the null hypothesis

A difficult task in permutation tests is strictly defining the null hypothesis under consideration. We focus on exchangeability as perhaps the most general and useful form of the null hypothesis in this setting. This hypothesis is defined as irrelevance of the labels in the form of an identical distribution being obtained under permutations. We formalize the concepts below.

Recall that *W_ij_* is the *l* dimensional feature vector of subject *i* on occasion *j* where *i* = 1*, · · ·*, *n* and *j* = 1, 2. Denote *W*_(*j*)_ as the *l × n* data matrix for occasion *j* = 1, 2 with columns *W*_1*j*_, · · ·, W_*nj*_. Let *W* = [*W*_(1)_*, W*_(2)_] be the *l ×* 2*n* combined data matrix with columns *W*_11_*, W*_21_*, · · ·, W_n_*_1_*, W*_12_*, W*_22_*, · · ·, W_n_*_2_. Let *W* = *w* be the observed data. Recall also, in matching with replacement, the best match for subject *i*’s occasion 1 image is *m_i_* = argmin*_r_ d*(*w_i_*_1_*, w_r_*_2_). In matching without replacement, the best match for subject *i*’s occasion 1 image is *m_i_*, where 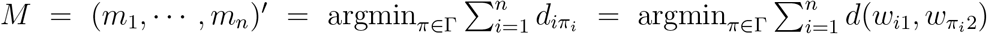, Γ is the collection of permutation vectors of (1*, · · ·, n*)^*′*^. In both scenarios, the test statistic is defined as 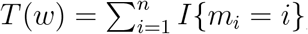, the number of correct matches.

The exchangeable null hypothesis, *H_E_*, is defined as the invariant distribution of test statistic when permuting the labels of occasion 2 images. That is,

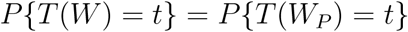

for all *t* ∈ {0*, · · ·, n*}, *P* ∈𝒫, where 𝒫 is the collection of *n×n* permutation matrices, *W_P_* = {*W*_(1)_*, W*_(2)_*P* } is the *n ×* 2 data matrix obtained after permuting occasion 2 labels.

### 3.2 Exact permutation tests

Following Hoeffding (1952), under *H_E_*, the permutation test can be executed to have an exact *α* type I error rate if a randomized test function is defined as:

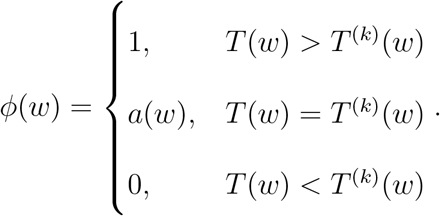

Here, *ϕ*(*w*) is the probability of rejecting the null given observation *W* = *w*. The variables, *T*^(*k*)^(*w*), for *k* = 1*, …, n*! is the ordered list of all permuted test statistics. The index *k* determines the closest quantile less than or equal to *α* of the permuted test statistics level, i.e. *k* = *n*! *–* ⌊*n*!*α*⌋ where ⌊*·*⌋ is the floor function. This is equivalently, the inverse, F̂^−1^(1 *− α*), of the distribution function of the permuted test statistics:

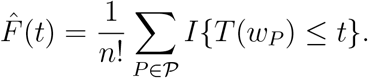

A randomized test with exact level *α* occurs if one rejects *H_E_* when *ϕ*(*w*) is 1, i.e. the test statistic lies strictly in the upper *α* area of the permutation distribution, fails to reject when *ϕ*(*w*) is 0, and rejects with probability *a*(*w*) otherwise. In the latter case, a uniform random variable is simulated and the test is rejected if it is less than *a*(*w*).

Hoeffding (1952) showed that *a*(*w*) defined as {*n*!*α − M*^+^(*w*)*}/M*^0^(*w*) yields an *α* level randomized test. Here, *M*^+^(*w*) and *M*^0^(*w*) are the counts of permuted statistics larger than or equal to *T*^(*k*)^, respectively. These are formally defined as: *M*^+^(*w*) = *|*{*j* ∈ {1*, · · ·, n*!} : *T*^(*j*)^(*w*) *> T*^(*k*)^(*w*)}*|* and *M*^0^(*w*) = *|{j* ∈{1*, · · ·, n*!} : *T*^(*j*)^(*w*) = *T*^(*k*)^(*w*)}*|* (see Appendix Section 1).

Since having an ancillary coin flip determine rejection is not desirable, the more conservative non-randomized test simply uses the non-randomized test function:

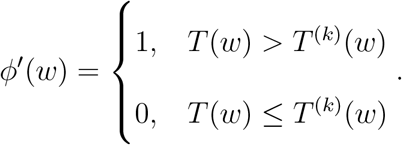

This yields a test with a type I error rate guaranteed to be less than *α*, though cannot yield an exact *α* level test, except in rare cases, such as when *n*!*α* is an integer.

Note that with the matrix representation we have *T* (*w*) = *tr*(*B*) as the total number of correct matches and hence *T* (*w_P_*) = *tr*(*BP*) = *tr*(*P B*) is the total number of correct matches after permuting occasion 2 labels according to some *P* ∈ 𝒫. Therefore an alternative expression for permutation distribution function is:

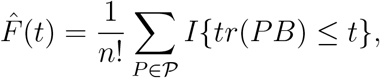

the CDF from the traces of all the row permutations of *B*.

Thus, the CDF arising from placing equal (discrete uniform) probability on all permutations is derived equivalently from permuting either the occasion 1 or occasion 2 labels.

### 3.3 Poisson approximation

#### 3.3.1 Matching without replacement

In matching without replacement, each occasion 1 image is matched to a distinct occasion 2 image. This implies each column and row of *B* sums to 1, as *B* is a permutation matrix, since the vector of matches is a permuted version of (1*, · · ·, n*)^*′*^ In this case, permuting occasion 1 labels and then calculating *tr*(Π*B*) is equivalent to shuffling a batch of ordered cards and counting the number of cards still in its original order, which follows Montmort’s matching distribution (Barton, 1958). Hence

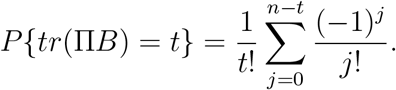

As *n* goes to infinity, for any fixed *t*, 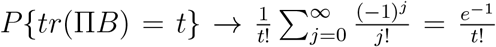 and *tr*(Π*B*) converges to a Poisson(1) distribution.

This is the distribution of correct matches under permutations, famously originally derived in a letter between Montmort and Nicolaus Bernoulli. This distribution and matching setting is often used in probability courses to illustrate the law of total probability. It is interesting to note that the Poisson approximation has an upper 95^*th*^ percentile of 3, 99^*th*^ percentile of 4 and 99.9^*th*^ percentile of 5. The relatively few matches need be made to reject this null hypothesis and that number is fairly static with *n*, since convergence occurs quite quickly. The reason the p-value is robust to large changes in *n* is because although the number of possible matches increases with *n*, the probability of a match decreases in a balanced way.

#### 3.3.2 Matching with replacement

Suppose we observed combined data matrix *W* = *w* and its representation matrix *B* in a matching with replacement process. Each occasion 1 image will be matched to exactly one occasion 2 image whereas some occasion 2 images may get matched multiple times and some occasion 2 images may not get matched at all. In this case the sum of any row of *B* will still be 1 but column sums of *B* can vary.

Without loss of generality, suppose only the column sums of first *k* columns of *B* are nonzero. Denote the column sums as *c*_1_*, · · ·, c_k_*. Then 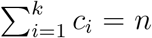. For *h* ⊂{1*, · · ·, k*}, denote the size of *h* as *|h|*. By the inclusion-exclusion formula we have (see Appendix Section 2)

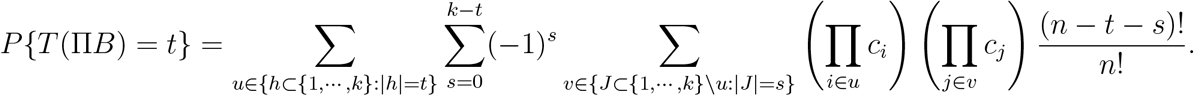

When *k* = *n* and *c*_1_ = *· · ·* = *c_n_* = 1, the distribution coincides with the matching without replacement distribution:

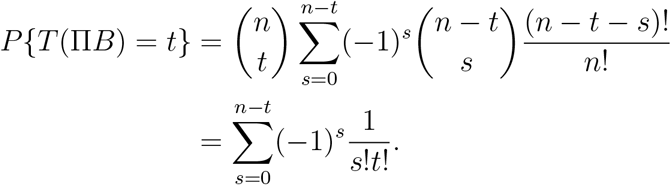

Via Stein-Chen’s method (see Appendix Section 3), the total variation between *T* (Π*B*) and a Poisson(1) for matching with replacement is:

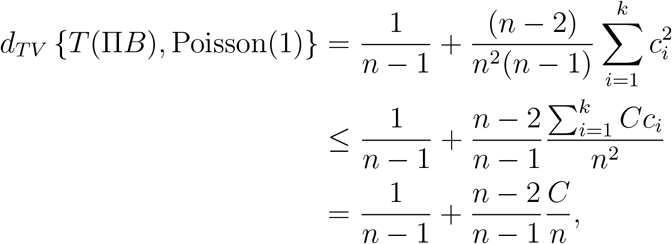

where *C* is the number of matches of the occasion 2 image with the most matches, that is, *C* = max_*i∈*{1*,···,k*}_ *c_i_*. Thus the permutation distribution will be approximated by a Poisson(1) if *C* is small and *n* is large. Specifically, 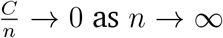 as *n → ∞* is sufficient for the distribution of *T* (Π*B*) to converge to a Poisson(1).

### 3.4 Power of the Poisson test

Consider an exchangeability permutation test for matching with replacement of level 0.05 approximated by a Poisson(1) distribution having test function:

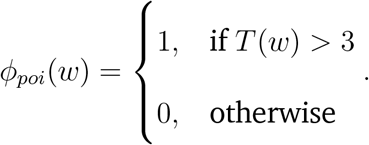

For *i, j* ∈ {1*, · · ·, n*} let the random variables *𝓑_ij_* = *I*{**m**_*i*_= *j*}, *𝓑_i_* = (*𝓑_i_*_1_*, · · ·, 𝓑_in_*)^*t*^ and matrix *𝓑* = (*𝓑*_1_*, · · ·, 𝓑_n_*) be unrealized version of B. Thus, for a realized observation, *W* = *w* we have *𝓑* = *B* and *T* (*w*) = *tr*(*B*). Assume *𝓑_ij_* follows a Bernoulli distribution with mean *p_ij_*. Then, for any sequence of distributions 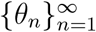 the power of the test is:

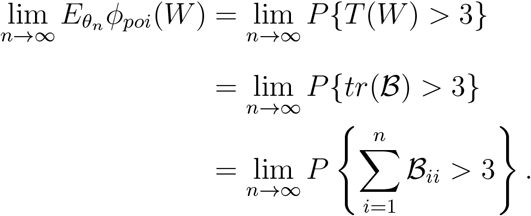

Consider an alternative hypothesis, *H_A_*_1_, where for all *θ_n_* we have that: (i) *B_ii_* (*i* = 1*, · · ·, n*) are iid with a Bernoulli distribution and a mean 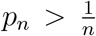 so that each subject will be more likely matched to themself than by chance, and 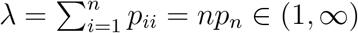. Then,

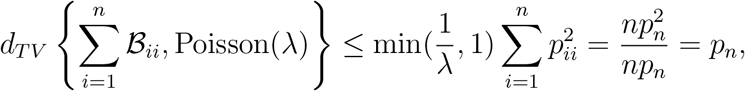

where *p_n_* = *λ/n →* 0. It follows that

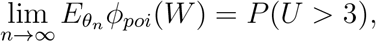

where *U ~* Poisson(*λ*).

Consider a more realistic alternative hypothesis, say *H_A_*_2_, where for all *θ_n_*: (i) 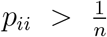 for all *i* = 1*, · · ·, n*, (ii) 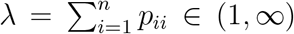 and (iii) *|cor*(*B_ii_, B_jj_*)*|* are small enough so that 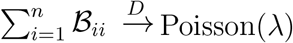 Poisson(*λ*). Such test will as well lead to a power of *P* (*U >* 3) where *U ~* Poisson(*λ*).

Figure 1 displays power, *P* (*U >* 3), against *λ*. Recall, *λ/n* can be interpreted as the average chance of getting a correct match. The test has a power greater than 80% when *λ* is roughly larger than 6. It also demonstrates a potential scenario of the test being under-powered, for example when subjects are guaranteed to be matched to themselves with a probability as large as three times of that by chance 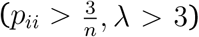. The test would end up with a power under 0.4, no matter how many subjects are recruited. However, unintended high power might occur if a higher *λ* is achieved due to the existence of twins, family or covariate structures. This is potentially problematic, as then the measure of reproducibility is highly sensitive to sample demographics and other factors that are generally not thought of as a component of reproducibility.

**Figure 1:**
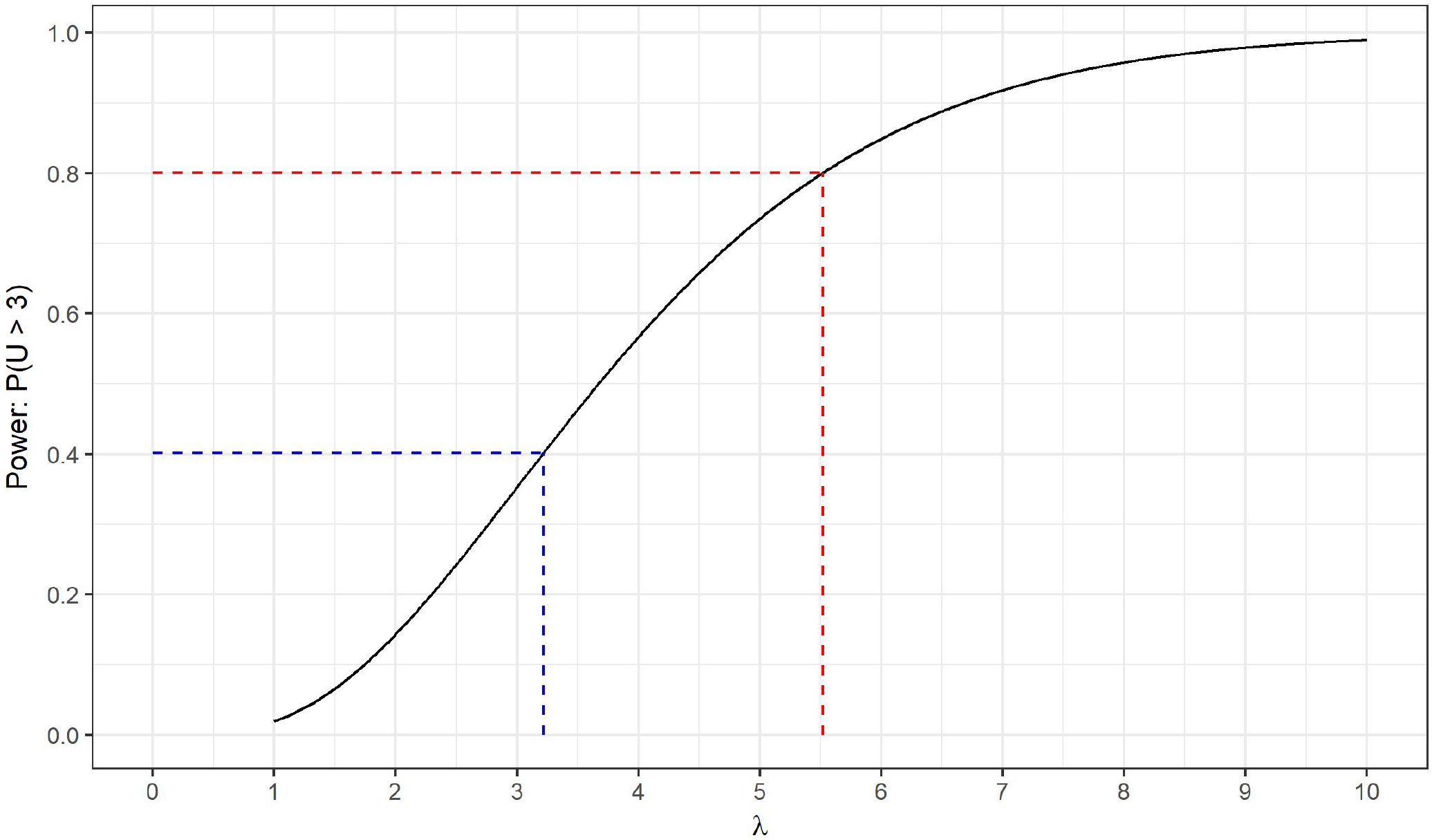
The power of the Poisson approximated permutation test with the alternative hypothesis *H_A_*_2_ (see Section 3.4) as *λ* changes.

### 3.5 Alternative justifications of the permutation test

Matching with replacement with common permutations represents the most popular form of permutation test for assessing reliability in fMRI. An often cited rationale behind this permutation scheme is to address potential correlations between subjects. As an example, consider if there are twins in the study. The null distribution still specifies that every second occasion match is equally likely, but that the twins are more likely to match to the same image. That is, for some pairs of subjects, under the null

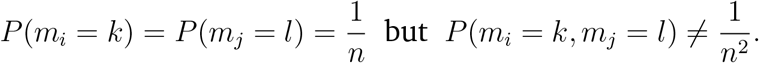

This permutation strategy addresses this potential correlation by conditioning on observed matching concordance. If any two subjects agree on a match, then will agree on all permutations and similarly any two subjects that disagree.

However, note that such tolerance of inter-subject correlation could be limited depending on the observed matches. For example, after permuting rows of the representation matrix, *B*, from a matching with replacement procedure, we have the covariance of two distinct subjects both getting correct matches to be:

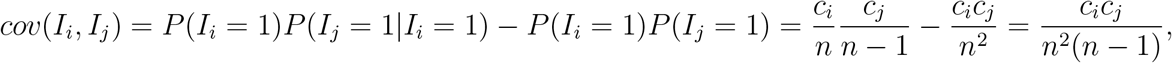

where *I_i_* denotes an indicator that subject *i* gets a correct match after permutation. Thus,

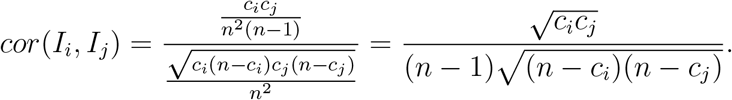

Therefore, a potentially small positive correlation is enforced by the permutation, no matter what the actual correlation is.

An alternative justification of this strategy lies in a conditioning argument. First, note that our permuted statistic satisfies: *tr*(*P B*) = *tr*(*BP*). Therefore, the null distribution is the same regardless of whether rows or columns are permuted. However, the row sums are all 1 and thus permuting the rows results in selecting a random table with fixed row and column sums. This is the same distribution obtained when considering generalizations of Fisher’s exact test of equality of the mulitinomial probabilities across rows, though the statistic in that case is usually a Chi-squared statistic or deviance, whereas we are simply considering the trace.

As is well known, this null distribution can be arrived at via conditioning on sufficient statistics. Consider a null hypothesis where the rows of *B* are assumed to be independent multinomial(*π*, 1) where *π* is an *n ×* 1 vector of probabilities. If one conditions on the sufficient statistics for *π*, which is the column sums (recall that the row sums are all 1), the resulting null distribution is then uniform on the space of tables satisfying the margins. The uniformity arises over the central hypergeometric distribution by virtue of all of the row margins being 1. This way of thinking is potentially useful for specification of the hypothesis being testing using this permutation scheme.

## 4 Numerical experiments

### 4.1 Poisson approximation of the null distribution

We will now demonstrate numerically how the permutation distribution under the null hypothesis *H_E_* is approximated by Poisson(1) in both matching without replacement and matching with replacement.

Consider *n* = 10, 25, 100, 500 samples each with measurements on visits *j* = 1, 2. The permutation distribution *F_tr_*_(Π*B*)_(*t*) is decided by the column sums of the representation matrix *B*. For MWOR, such column sums are all 1’s. For MWR, we simulated a matrix *B* for each sample size *n* following the rule that each subject would get matched to all subjects with equal probabilities, in which case, *C*, the number of maximum matches a subject could get would increase with a slow rate 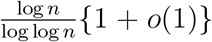 so that *C/n →* 0 with probability 1 *− o*(1) (Raab and Steger, 1998). Both data generating distributions satisfied the exchangeability assumption *H_E_*.

Within each iteration, we randomly permuted the columns of the simulated *B* matrices once and then recorded the observed traces. The total number of iterations was 1,000. We plotted the simulated permutation distributions for sample sizes *n* = 10, 25, 100, 500 and for MWOR and MWR with a comparison against the density of Poisson(1) distribution in Figure 2.

**Figure 2:**
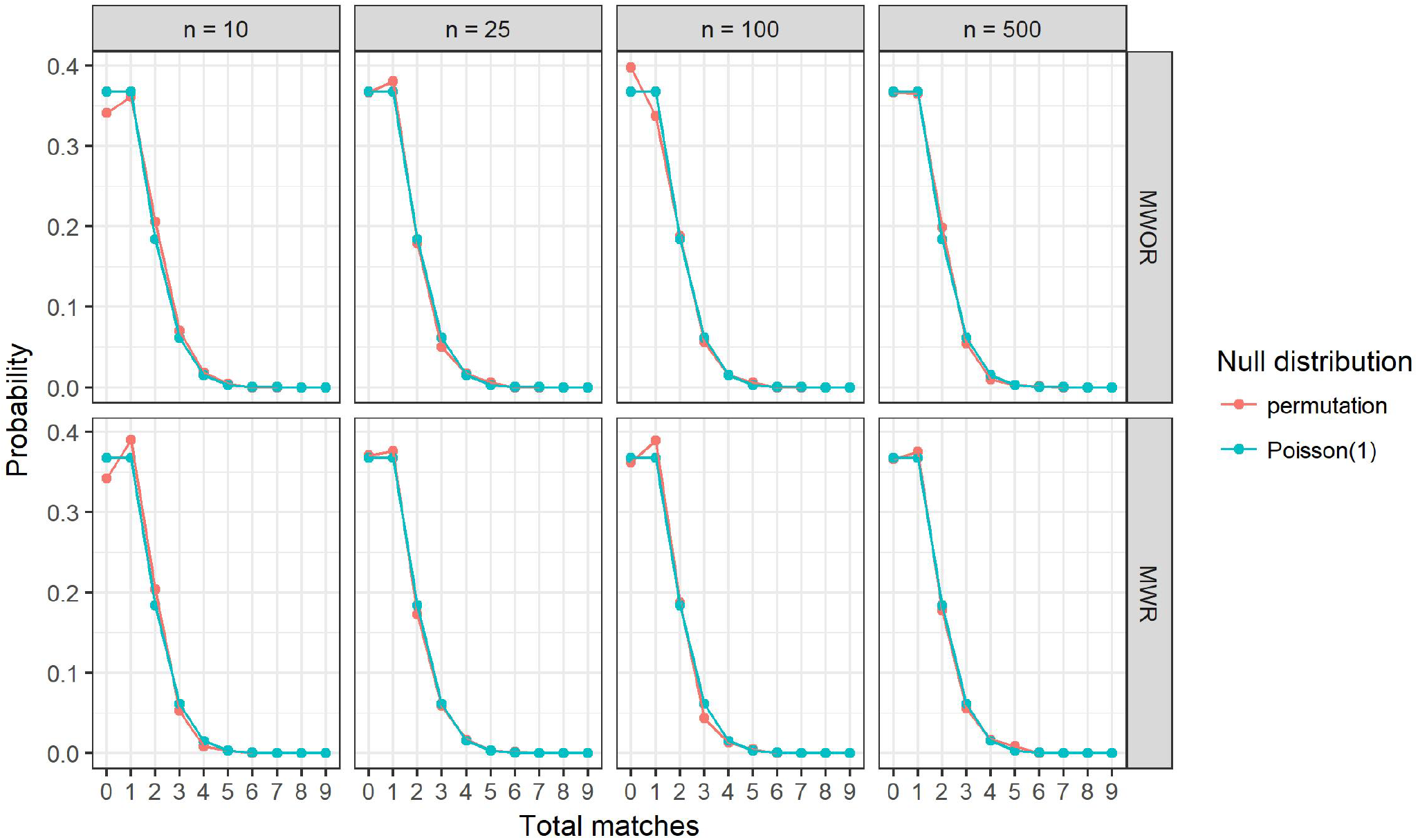
The simulated permutation distributions for MWOR and MWR after 1,000 iterations under the null hypothesis *H_E_* with sample sizes *n* = 10, 25, 100, 500 compared with Poisson(1) distribution (see Section 4.1). The permutation distributions are in red and the Poisson(1) distributions are in blue.

### 4.2 Poisson approximation on BLSA and HCP datasets

For the Baltimore Longitudinal Study of Aging (BLSA) dataset, 277 older participants (151 females, age 55 to 96) were included from the neuroimaging substudy of the BLSA (Resnick et al., 2000) who got rs-fMRI scans on multiple visits. The first and last available images of each participant, 554 scans in total were used in the dataset for matching. The time intervals between the first and last available images of the subjects ranged from 310 days to 1, 799 days.

For the Human Connectome Project (HCP) dataset, 466 participants (273 females, age 22 to 36), each with two separated resting state fMRI sessions on consecutive Day 1 and Day 2, were included from the HCP (Van Essen et al., 2013) S500 release. Preprocessing was conducted following the minimal preprocessing pipelines (Glasser et al., 2013). For each participant, the rs-fMRI scan with the left-to-right phase encoding direction in each session was used so that we had 932 scans in total for matching.

On both datasets, the atlas with 268 nodes partitioned into eight networks defined with the Shen’s functional parcellation method on the independent health controls (Shen et al., 2013; Finn et al., 2015) was applied to each rs-fMRI image. The feature vector *W_ij_* for subject *i* on the first (*j* = 1) or the last (*j* = 2) visit was taken as the upper triangular of the Pearson correlation (z transformed) matrix calculated for all the nodes using their time series during the corresponding scan. The distance, *d*(*·, ·*), was defined as one minus the Pearson correlation between the two feature vectors.

Within each iteration, we took random subsamples with numbers of subjects *n* = 10, 25, 100, and all subjects for both datasets. Matching with replacement was conducted. The test statistic was the number of total matches or the trace of the representation matrix *B*. A permutation test and a Poisson test at level 0.05 then followed. For each of the eight scenarios (which are the combinations of the four subsample sizes and the two datasets), 1,000 permutation p-values and 1,000 Poisson p-values were obtained after 1,000 iterations.

The Poisson and permutation tests agreed on rejection of the null in all but five iterations, all of which were at size *n* = 10 from the BLSA dataset. When sample size *n* = 25, 100, or all subjects, reported p-values and distances between the two types were less than 0.001 in all iterations (see Table 3).

**Table 3:** Illustrating the accuracy of the Poisson(1) approximations. Distribution of the distances between the Poisson and permutation p-values in Section 4.2 are given. Tests were conducted on random subsamples with sizes *n* = 10, 25, 100, an all of the subjects from the BLSA or the HCP datasets. Matching with replacement was conducted. The total number of iterations was 1,000 for all scenarios.

|  | n = 10 |  | n = 25 |  | n = 100 |  | full data |  |
| --- | --- | --- | --- | --- | --- | --- | --- | --- |
|  | BLSA | HCP | BLSA | HCP | BLSA | HCP | BLSA | HCP |
| [0,0.001) | 899 | 995 | 1000 | 1000 | 1000 | 1000 | 1000 | 1000 |
| [0.001,0.01) | 91 | 5 | 0 | 0 | 0 | 0 | 0 | 0 |
| [0.01,0.1) | 10 | 0 | 0 | 0 | 0 | 0 | 0 | 0 |

**Table 4:**
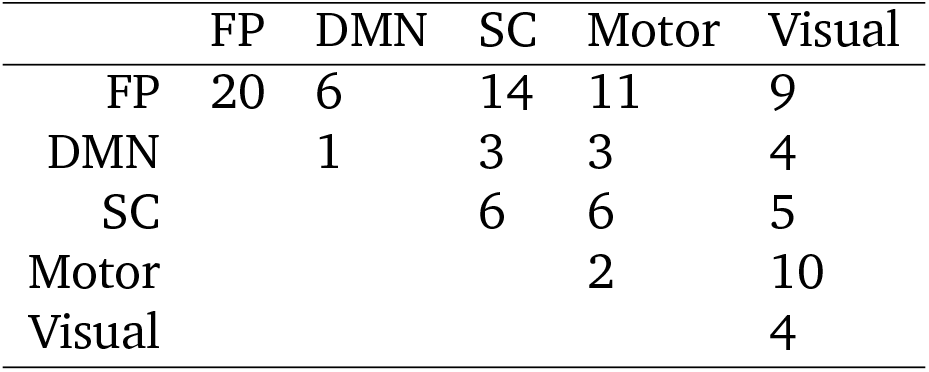
Numbers of the identifying pairs between the five combined networks on the HCP dataset (see Section 4.8). The pairs of nodes were selected by the Poisson approximated permutation test on the total matches from matching with replacement using only the z transformed correlations between each single pair.

**Table 5:** Identification rates (in %) from matching with replacement using only the z transformed correlations of the pairs between the five combined networks on the HCP dataset (see Section 4.8).

|  | FP | DMN | SC | Motor | Visual |
| --- | --- | --- | --- | --- | --- |
| FP | 90.6 | 80.7 | 70.0 | 61.2 | 66.7 |
| DMN |  | 50.0 | 56.7 | 33.7 | 45.5 |
| SC |  |  | 44.6 | 45.7 | 53.6 |
| Motor |  |  |  | 42.3 | 45.7 |
| Visual |  |  |  |  | 58.8 |

### 4.3 Sensitivity to uninteresting directions of the alternative given existence of clustering structures

In this section, it is demonstrated that the fingerprinting permutation test may produce undesired significance when clustering structures exist.

Consider a one dimensional measurement, *W_ij_*, with a categorical covariate, *Y_i_*, defined as follows for subject *i* on visit *j*, *i* = 1*, · · ·, n* = 300, *j* = 1, 2. The *n* subjects are partitioned by *K* clusters, each with size 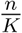. Let *Y_i_* = *k* if subject *i* belongs to the *k*-th cluster and *W_ij_* = *Y_i_ ·* 100 + *r_ij_* where *r_ij_*’s are iid variables uniformly distributed on [*−*50, 50]. Therefore subjects are exchangeable within clusters with a probability of matching to themselves as 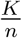.

According to the power analysis for alternative hypothesis in the previous section, a permutation test of level 0.05 with Poisson approximation on such data will lead to a power of *P* (*U >* 3) where *U ~* Poisson(*K*).

We simulated the distribution of the statistic, the number of matches *T*, with *B* = 1,000 iterations for *K* = 2, 4, 6, 10. Within each iteration, subjects were partitioned into *K* clusters, *W_ij_, r_ij_*’s were then simulated and matching with replacement was conducted accordingly. We plotted the simulated distribution of *T* for different clustering settings (Figure 3). One will observe that the test could reject the null hypothesis *H_E_* with probabilities as high as 80% or 99% with as few as *K* = 6 or *K* = 10 clusters.

**Figure 3:**
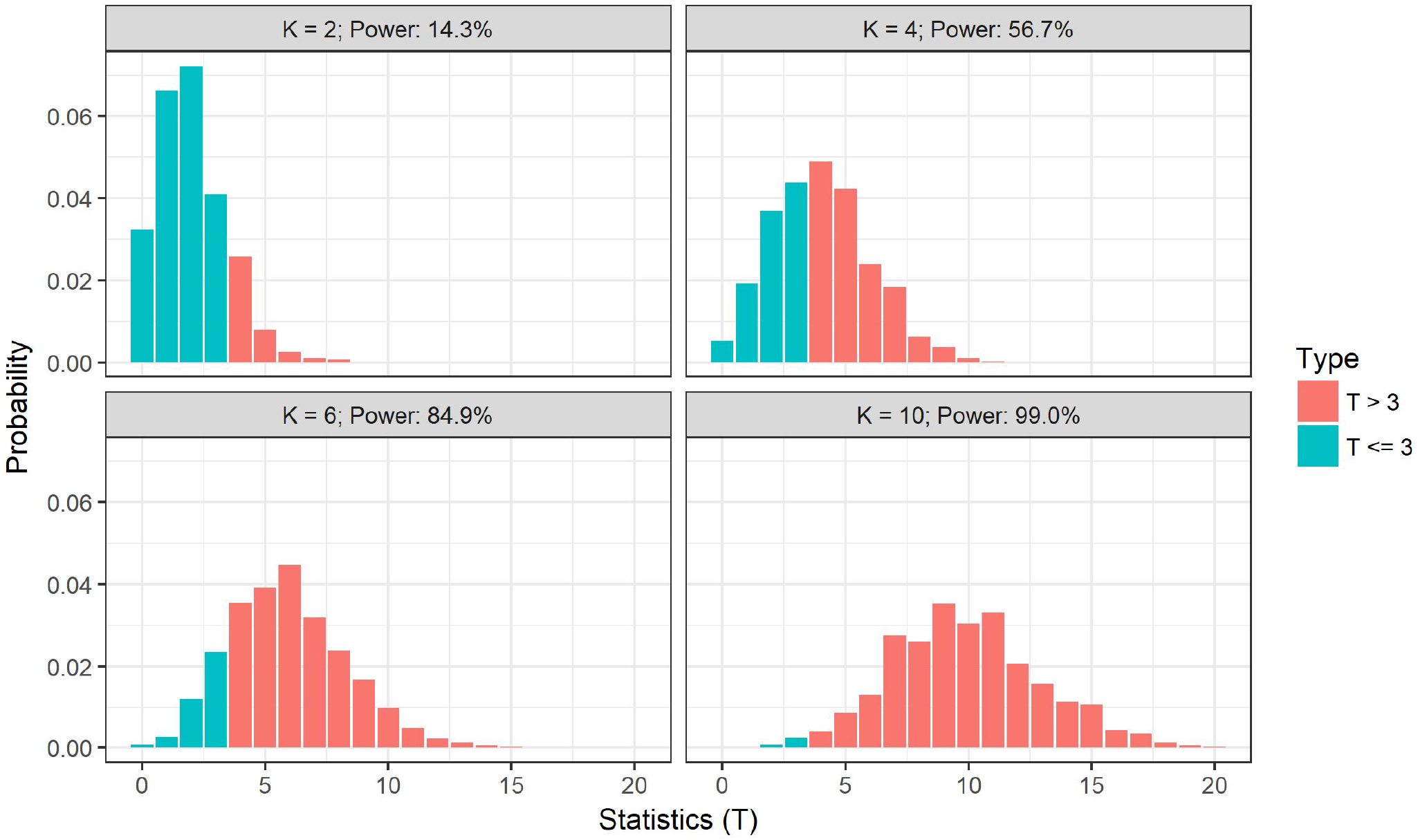
Simulated distributions of the numbers of matches (*T*) with the clustering settings in Section 4.3. *K* = 2, 4, 6, 10 were the numbers of clusters. Matching with replacement was conducted. For a Poisson test at level 0.05 it would reject the *H_E_* when *T >* 3, which was colored in red, otherwise in blue. Powers of the Poisson tests calculated from Section 3.4 were also labeled for all scenarios.

However we argue that such high power is hardly desirable due to the fact that observations are iid within clusters for all subjects and visits. This demonstrates that one might reject the null simply because of the demographics of the sample or clustering factors not generally thought of as related to reproducibility. In such settings, the fingerprint test can strongly reject the null, despite the fact that there contains nothing identifying in the measurement other than covariate information. To relate this to a practical setting, an fMRI study with varying ages will be more likely to reject than the same measure in a study with constant ages.

### 4.4 Matching on a contaminated HCP dataset

We will demonstrate the issue of being over-sensitive or over-powered by a matching example on the HCP dataset where we deliberately contaminate different proportions of the scans on one occasion with irrelevant scans from the BLSA dataset.

Recall our HCP dataset with 466 participants and 932 scans. Within each iteration, we took random subsamples with numbers of subjects n = 25 or 100. Then we randomly chose 50%, 75%, 90%, 95%, or 100% (rounded to the closest integer) of the occasion 1 scans from the selected subsamples, where we replaced them with scans randomly chosen (without replacement) from the 554 BLSA scans. Matching with replacement was conducted. The permutation test and the Poisson test at level 0.05 then followed for all 1,000 iterations.

According to the Bland-Altman plots (Bland and Altman, 1986, 1999) (Figure 4), the two types of tests simultaneously rejected the null during 42.5% of the iterations with contamination ratio 90% when *n* = 25, and did so during 88.2% of the iterations with contamination ratio 95% when *n* = 100. Thus, one rejects the null hypothesis with high probabilities, even if over 90% of the scans on a visit are contaminated with irrelevant data.

**Figure 4:**
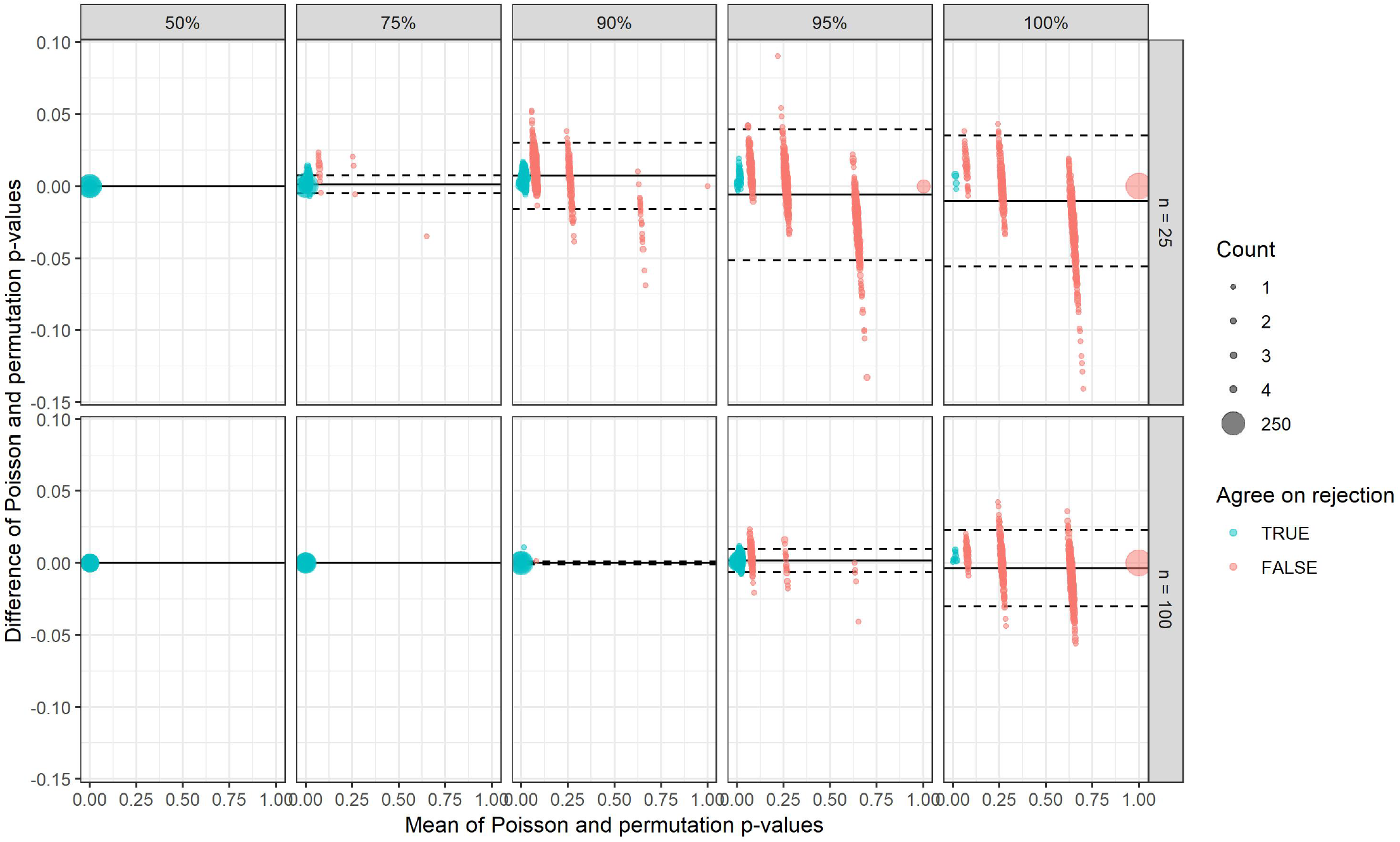
Bland-Altman plots for Poisson and permutation p-values in Section 4.4. Random subsamples of sizes *n* = 25 or 100 were taken from the HCP dataset during each iteration. 50%, 75%, 90%, 95%, or 100% (rounded to the closest integer) of the occasion 1 scans were replaced with randomly selected BLSA scans. Matching with replacement was conducted. The total number of iterations was 1,000 for all scenarios. Differences were plotted against the mean; 95% limits of agreement were plotted as dotted lines; mean levels were plotted as solid lines; overlapping dots were sized according to the counts. Points are color-coded as red if the two types of tests agreed on rejection of the null, otherwise as blue.

### 4.5 Matching for comparing connectome similarities between twins or non-twin siblings

Our HCP dataset included 53 families with monozygotic (MZ) twins and other 24 families with dizygotic (DZ) twins, all verified by genotyping. There were another 68 families with genotyping data available that had at least two siblings but no twins (NotTwin), which added up to 157 non-twin siblings.

Within each iteration, from each of the three types of families above (MZ, DZ or NotTwin), we randomly selected 20 families. Then from each of the selected families, we randomly chose an ordered pair of twins (for families with MZ and DZ twins) or non-twin siblings (for families with no twins but at least two siblings and with genotyping data available). We also randomly selected 20 ordered pairs of subjects from all the 466 participants (labeled Random).

For each selected ordered pairs, we took the measurement of the first experiment session for the first subject and that of the second experiment session for the second subject. Then for each of the four scenarios (MZ, DZ, NotTwin and Random), we had two groups of 20 measurements from totally distinct subjects.

If different levels of similarities between siblings existed, then the distributions of the total number of matches for siblings could diverge not only from that when siblings were no closer than random people and the exchangeability assumption held, i.e. a Poisson(1) distribution, but between those of different sibling types as well.

After 1,000 iterations the empirical distributions were plotted (Figure 5). An empirical distribution of 1,000 iid Poisson(1) samples was also plotted as comparison. It shows that the total number of matches followed a similar distribution for completely random samples and the Poisson(1) samples, with the proportions of rejecting the null at level 5% in the Poisson tests being 1.1% and 1.4% respectively, which coincided with the probability 1.9% of being greater than 3 for Poisson(1). We also observed similar distributions for DZ twins and non-twin (NotTwin) siblings, with the proportions of rejecting the null at level 5% being 54.1% and 54.2% respectively. For MZ samples the numbers of matches were greater than 3 in all iterations. These results could also be seen as supportive evidence in terms of the brain connectivity for the genetic assumption that MZ twins having greater similarity than DZ twins or non-twin siblings, which were all closer than random pairings.

**Figure 5:**
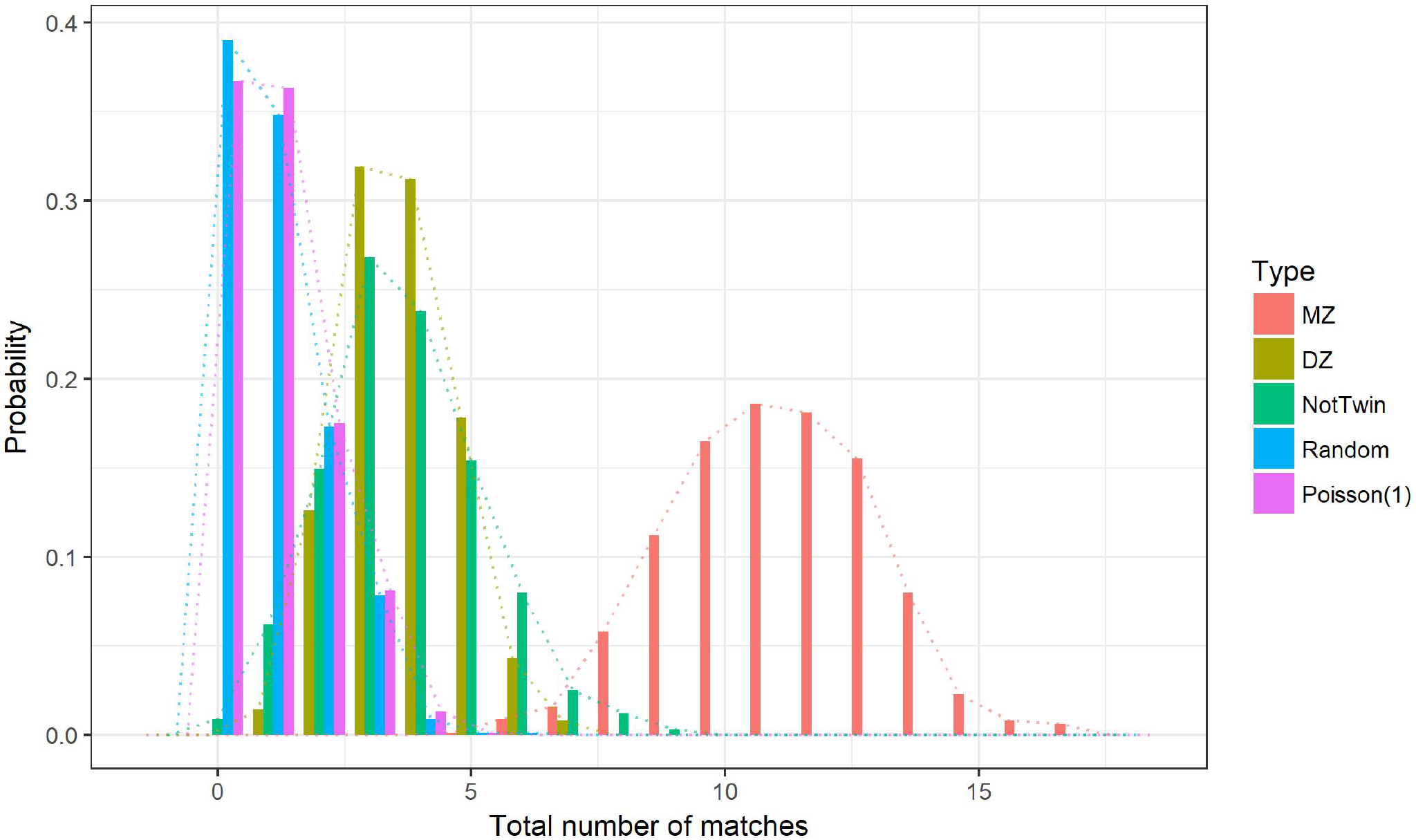
The simulated distributions of the total number of matches when matching two groups of distinct people (each of size 20) from the HCP dataset. For each person selected in the first group, there was another monozygotic twin/dizygotic twin/non-twin sibling/random person in the second group for the MZ/DZ/NotTwin/Random scenarios, respectively (see Section 4.5). Matching with replacement was conducted. The empirical distribution of a Poisson(1) random variable after 1,000 iterations is also plotted as comparison.

Such matching experiments between distinct subjects demonstrated how the fingerprint test when specially designed can serve as a test for the existence of similarity among people with certain social or genetic relations. According to the experiment results, the power of such a test could be relatively low (around 50% for the level of similarity between DZ twins or non-twin siblings) or very high (close to 100% for the level of similarity between MZ twins) for brain connectivity measurements depending on the (usually unspecified) alternative hypothesis. The empirical distributions of the test statistic demonstrated a way of comparing the levels of brain connectome similarities for different genetic or social relations.

### 4.6 Covariates associated with matching in BLSA and HCP datasets

On the HCP dataset, matching with replacement on the 466 participants resulted in 350 people (75.11%) getting matched to themselves. Let 1 represent that a subject got correctly matched and 0 otherwise. Using a logistic regression model, we regressed the matches against demographic covariates, including years of education, age, sex, race (having levels “Asian/Native Hawaiian/Other Pacific Islander”, “Black or African American”, “White”, “More than one” and “Unknown or Not Reported”; “Asian/Native Hawaiian/Other Pacific Islander” as the baseline), income and whether the participant is still in school. Two variables were marginally interesting: age with estimated odds ratio 1.06, 90% CI [1.01, 1.12], Wald z statistic 1.80 and p-value 0.073; the race category for black or African American, having an estimated odds ratio 0.15, 90% CI [0.02, 0.94], Wald z statistic *−*1.70 and p-value 0.088. Though these variables show weak evidence for associations with matching, recall that the ages, ranging from 22 to 36 on the HCP dataset, were all health healthy and younger.

We further investigated if any similarity in terms of resting state connectivity existed among people with the same age and race category. Within each iteration, from each of the 208 families we randomly selected one subject so that no sibling structure existed. We then partitioned the 208 subjects by age and race categories. We randomly chose 20 combinations of age and race categories that contained more than one subject in the 208 samples. From subjects with each of the selected age and race combination, we then randomly chose an ordered pair of subjects. For the first subject of a pair we took the measurement of the first experiment session and for the second subject we took that of the second experiment session. We then conducted matching with replacement on the two groups of 20 measurements, now having totally distinct participants on the two session. After 1,000 iterations the empirical distribution was plotted for the total matches with an empirical distribution from the previous iid Poisson(1) samples as comparison. From Figure 6 a slight right shift from the Poisson(1) was observed for the age and race matched simulated samples with a proportion of rejecting the null at level 0.05 in the Poisson tests being 6.3%, which was larger than that in the Poisson(1) samples as 1.4% and probability 1.9% of being greater than 3 for Poisson(1) distribution; these were substantially smaller than those in the dizygotic twins (54.1%) or non-twin siblings (54.2%).

**Figure 6:**
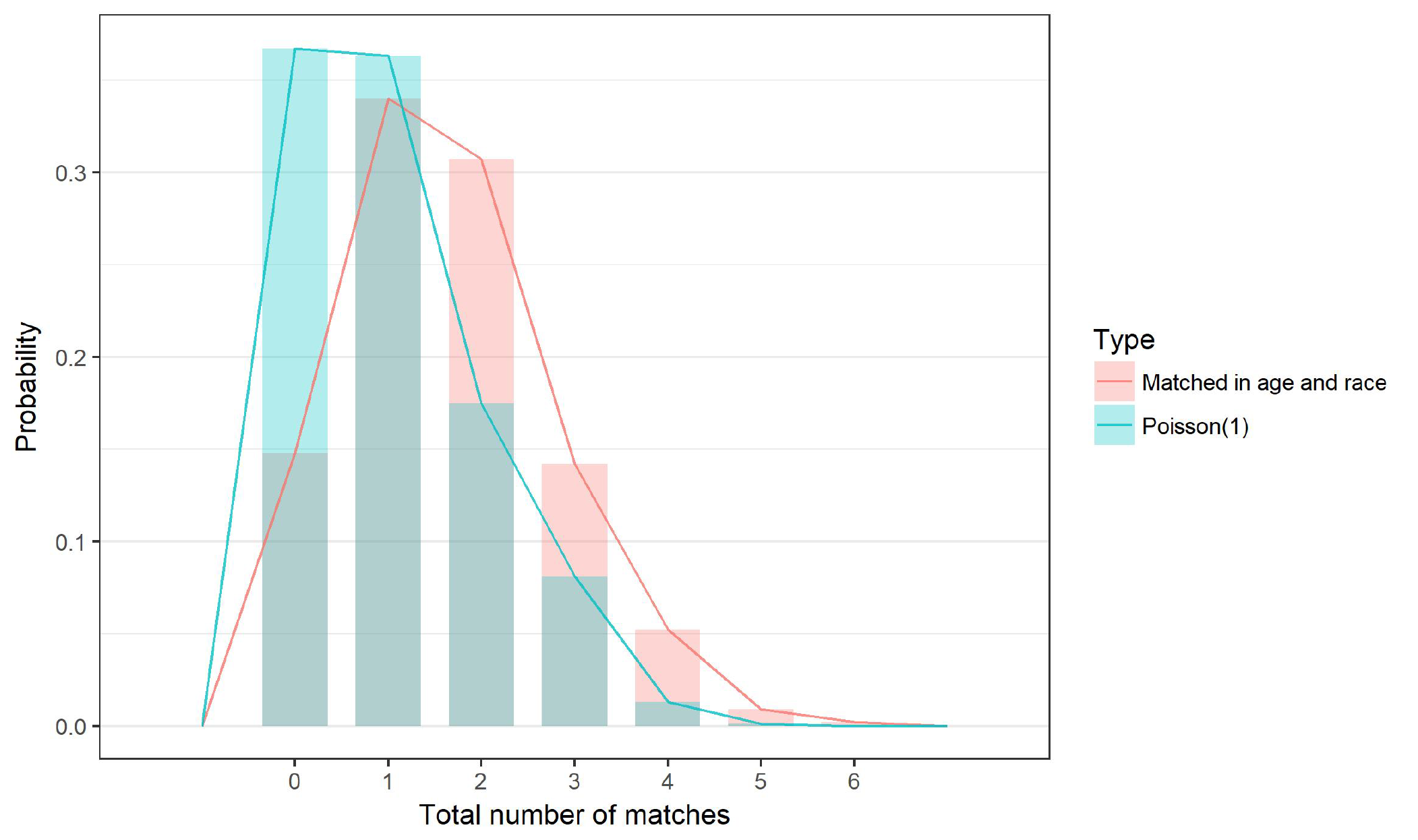
The simulated distribution of the total number of matches when matching two groups of distinct people (each of size 20) who were randomly selected from different families and were matched in age and race in the HCP dataset (see Section 4.6). Matching with replacement was conducted. The empirical distribution of a Poisson(1) random variable after 1,000 iterations was also plotted as comparison.

On the BLSA dataset, matching with replacement on the 277 participants resulted in 110 people (39.71%) getting matched to themselves. We again applied a logistic regression model for match status against demographic covariates including: years of education, age on the first scan, sex, race (with 3 levels as “Black or African American”, “White” and “Other”) and the time interval between the two scans in days. At level 0.05 two variables were potentially related to matching status: years of education with estimated odds ratio 1.12, 95% CI [1.00, 1.25], Wald z statistic 2.02 and p-value of 0.043; sex with estimated odds ratio 1.80, 95% CI [1.09, 2.98], Wald z statistic 2.28 and p-value of 0.023. It is important to emphasize the ages in the BLSA dataset ranged from 55 to 96 (far different from that of the HCP dataset) and there was a much larger time span between scans.

We could visualize such observation by comparing matching on random subsamples from the whole dataset and matching on those from a subgroup of participants whose years of education were higher than the dataset average and sex codes were 1’s (Figure 7). Such subgroup included 69 participants out of the total 277 participants. Within each iteration, we took 20 random samples from such subgroup and from the whole dataset, respectively. Matching with replacement was then performed among their first and second scans. After 1,000 iterations, total numbers of matches were plotted for subsamples from the subgroup and those from the whole population. All observed numbers of matches exceeded 3 so we ended up with a proportion 100% of rejecting the exchangeability assumption in both scenarios. However the mean of the simulated numbers of matches in the subgroup was 13.73 (with SD 1.84) which was greater than that in the whole dataset (mean 12.74, SD 2.07).

**Figure 7:**
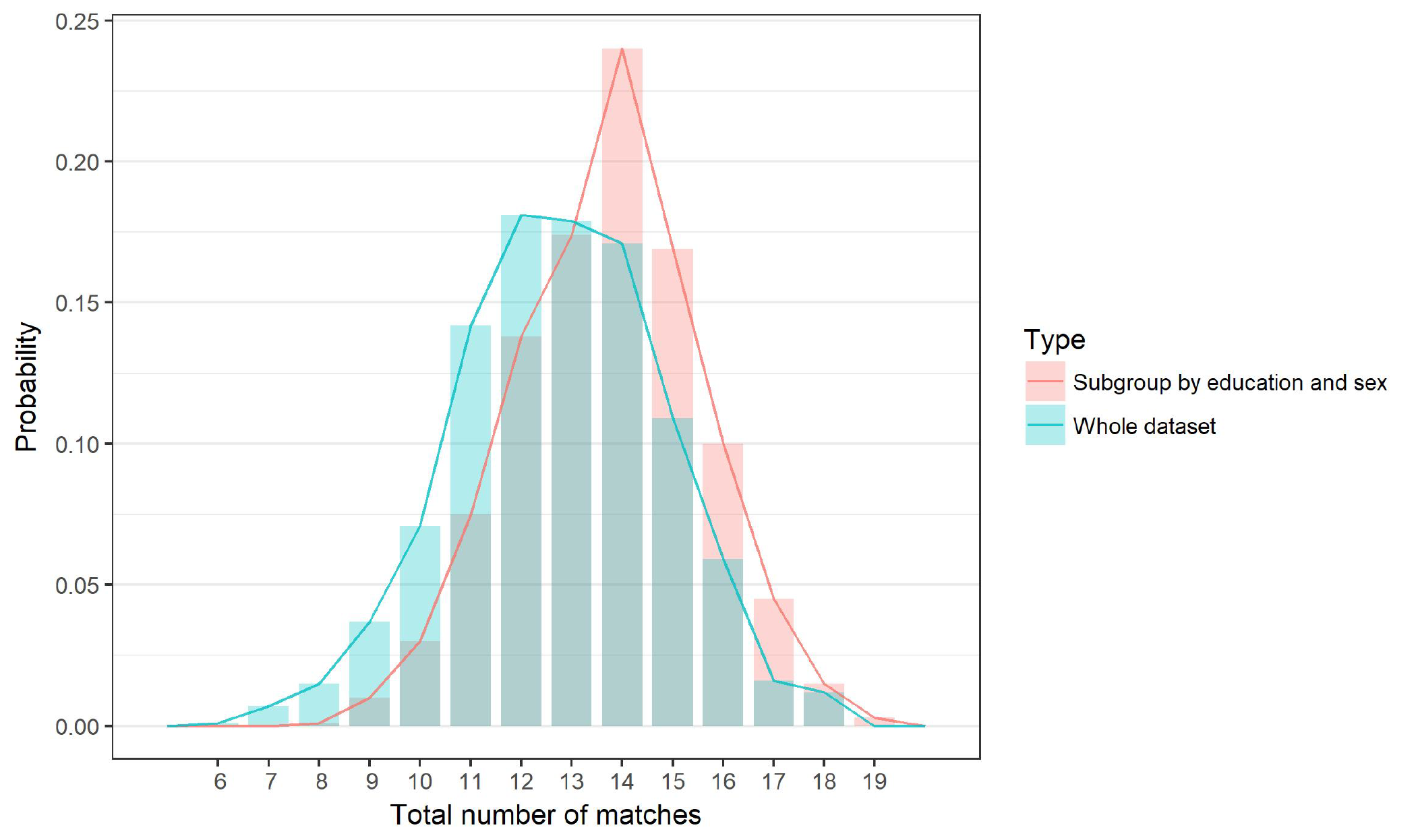
The simulated distribution of the total number of matches when matching with replacement on random subsamples of size 20 from the whole BLSA dataset or from the subgroup of participants whose years of education were higher than the dataset average and sex codes were 1’s (see Section 4.6).

### 4.7 Comparison of matching on the BLSA and the HCP datasets

The identification accuracy on the BLSA dataset was relatively low (39.71%) compared to that on the HCP dataset (75.11%). Such a drop in identification accuracy might partially be explained by the difference in the temporal resolutions and scan durations (HCP: TR/TE 720/33.1 ms, frames per run 1200, run duration 14 min 33 sec; BLSA: TR/TE 2000/30 ms, frames per run 180, run duration 6 min). A similar drop in identification accuracy has been reported previously when comparing matching with replacement on the data with more standard quality to that on the HCP data (Waller et al., 2017). It is also possible that the long time span between scans and more advanced ages of the BLSA participants is evidence of actual biological aging so that subjects are less like their previous selves, thus correctly having fewer matches.

However, at least two factors can make such comparison of identification rates suboptimal. First, the sample size could have an impact on the identification accuracy (Waller et al., 2017) and sample size differences exist (HCP dataset: 466; BLSA dataset: 277). Secondly, existing family structures such as the MZ twins could produce many matches, even between two group of totally distinct participants (see Section 4.5).

### 4.8 Brain maps of identifying pairs of nodes by network

Consider evaluating how well a single pair of nodes can identify people by conducting matching with replacement with only the single inter-node z-transformed correlations. Since the measurements are one dimensional we use the absolute difference as distance and randomly choose a match when ties appear. We use the Poisson approximation to the number of matches. An FDR adjustment follows for multiple testing. The Poisson approximation is useful in this setting, as the number of matching experiments grows with the order of the number of nodes squared.

On the HCP dataset using the sample of 466, 106 identifying pairs of nodes were discovered out of 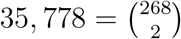 pairs (268 nodes). The total matches on those identifying pairs ranged from 7 to 10.

For simplicity, we combined the eight networks into five and then counted the identifying pairs between the following five combined networks: FP (the combination of Medial Frontal and Frontoparietal networks), DMN (Default mode network), SC (Subcortical-cerebellum network), Motor (Motor network) and Visual (the combination of Visual I, Visual II and Visual Association networks). FP was the network with most identifying pairs (20).

We further conducted matching with replacement only using the pairs between any two selected networks. It led to similar results that the identification rate on FP was the highest (90.6%). The 20 identifying pairs within the FP network are visualized (see Figure 8) on the ICBM 152 template brain (Mazziotta et al., 2001) with the **rgl** and **misc3d** packages in **R** (Adler et al., 2018; Feng et al., 2008; Muschelli et al., 2014).

**Figure 8:**
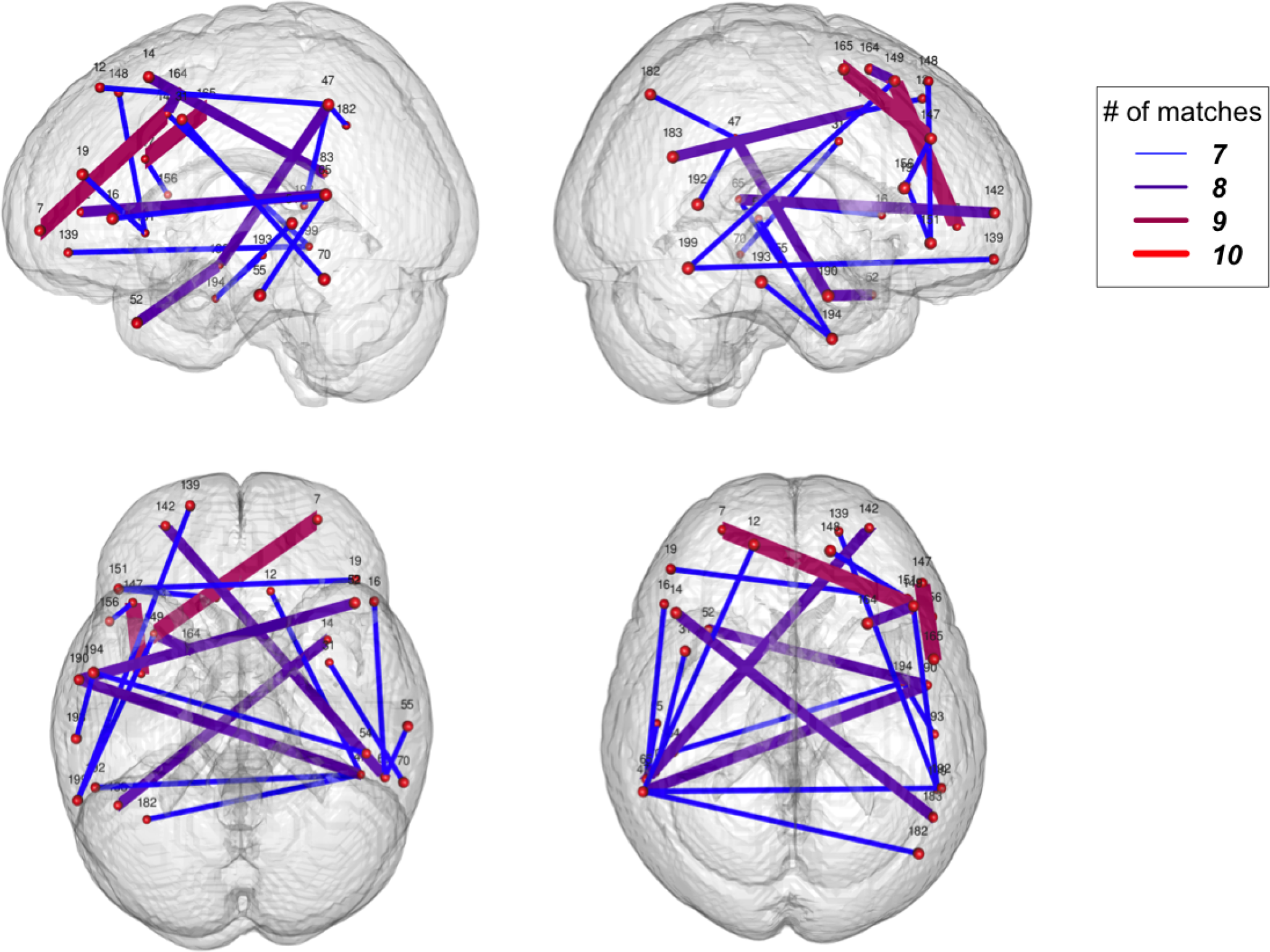
The 20 identifying pairs of nodes within the FP network visualized on the ICBM 152 brain template (see Section 4.8). Nodes were labeled by their orders on the atlas and were plotted at the center. Pairs of nodes were colored from blue to red depending on the number of matches when matching with replacement was conducted with only the z transformed correlations between each single pair.

The matching performance over individual nodes mirrors neuroscientific intuition that frontal networks are more idiosyncratic and personal, while motor and visual networks are more common across individuals.

## 5 Discussion

In this manuscript we considered matching permutation tests for so-called fMRI fingerprinting. We found that, regardless of the permutation strategy, the tests results in a Poisson(1) null distribution for the number of correct matches. When matching with replacement, the maximum number of matches for a subject must go to 0 with *n* for convergence. In matching without replacement, Montmort’s famous result also implies a Poisson(1) distribution. Finally, (not discussed) if one were to permute after each match with replacement, the number of correct matches would be Binomial(*n,* 1*/n*), which clearly limits to a Poisson(1) as well.

Thus, one can compare the number of matches to the relevant upper quantile of a Poisson(1) without further computing. This is particularly useful for studies of individual brain locations, or pairs of locations. In these settings, the lack of need for calculating a permutation based null distribution dramatically reduces computing time. In addition, the high power of the test mitigates the need for elaborate multiple comparison procedures and simpler more conservative variations would likely suffice.

While nearly any reasonable permutation and matching strategy yields a Poisson(1) null distribution for the number of correct matches, there are differences between the strategies. For example, matching with replacement yields a different answer whether occasion 1 or 2 is used as the reference group. In addition, poor matching without replacement strategies can be dependent on the original subject ordering. Matching with replacement more easily generalizes to multiple measurements per subject.

The exchangeability test was seen to be very highly powered and sensitive to assumptions towards a greater propensity to reject. Most notably, any correlation of the measurement with a demographic or clustering variable will aid in matching. This is intuitive. If one had pairs of outfits from several people and had to match them up in the absence of the owners, the task would be much harder if everyone was the same size, gender, etc. This has implications for the use of fingerprinting as a measure of reproducibility. For example, it is well known that resting state fMRI data changes with age. For the same experimental protocol measures of reproducibility would change depending on the age variation of the study subjects.

In addition, it is not clear, even in the absence of covariate or clustering variables, that matching performance is a reasonable estimate of reproducibility. At the minimum, it must be combined with other metrics (such as the I2C2 Shou et al., 2013) and a study of matching performance and its associations. We suggest the use of logistic regression on whether or not subjects were correctly matched for this task.

Subject identification is also an incomplete measure of the performance of a metric. It is worth remembering that ones actual fingerprint itself is a very good identifier, but is otherwise biologically meaningless, whereas gender, sex, medication usage, etc. are all poor subject identifiers but scientifically useful.

The data analysis yielded several interesting findings. The percentage of correct matches varied quite a bit between the two studies. This is sensible, as the HCP data included technical, one day separated, replicates, with a narrow age range of healthy subjects and long scanning sessions, hence well measured resting state data, collected in part to study resting state reproducibility and narrow down optimal acquistion protocols. In contrast, with a lower percentage of correct matches, the BLSA data is longitudinal, with a year or more between scans, considering an age range where resting state phenotypes may be longitudinally changing from normal aging and early stage disease and the resting state data was acquired using a shorter protocol as part of a larger battery of scans to study many facets of brain aging.

The HCP data included twins and it interesting that matching performance followed the appropriate order (from best performance): self, monozygotic twin, dizygotic twin, non-twin sibling and stranger. Among the basic demographics, age, education and race showed some association with matching performance. Various numeric experiments showed that one can obtain a more significant result by making the distribution of the significant demographics more variable, even when matching to strangers.

The final analysis considered all pairs of regions separately. It was primarily frontal cortical regions that were the most fingerprint-like (i.e. idiosyncratic). This mirrors both intuition and general results in this area. Intuition would suggest, for example, that intra-motor or intra-visual, connections would be similar across a collection of typical subjects simply because of the consistency of motor and visual function.

For future research, it is perhaps worthwhile considering ranking rather than matching. The current style of analysis treats a person being their own second best match identically to being the worst match. A rank sum styled test called “discriminability” (Vogelstein et al., 2015; Airan et al., 2016) avoids this complication, but has a harder null distribution to consider.

